# Reward and expectancy effects on neural signals of motor preparation and execution

**DOI:** 10.1101/2021.07.02.450521

**Authors:** Xing-Jie Chen, Berry van den Berg, Youngbin Kwak

## Abstract

The prospect of rewards can have strong modulatory effects on response preparation. Importantly, selection and execution of movements in real life happens under an environment characterized by uncertainty and dynamic changes. The current study investigated how the brain’s motor system adapts to the dynamic changes in the environment in pursuit of rewards. In addition, we studied how the prefrontal cognitive control system contributes in this adaptive control of motor behavior. To this end, we tested the effect of rewards and expectancy on the hallmark neural signals that reflect activity in motor and prefrontal systems, the lateralized readiness potential (LRP) and the mediofrontal (mPFC) theta oscillations, while participants performed an expected and unexpected action to retrieve rewards. To better capture the dynamic changes in neural processes represented in the LRP waveform, we decomposed the LRP into the preparation (LRP_prep_) and execution (LRP_exec_) components. The overall pattern of LRP_prep_ and LRP_exec_ confirmed that they each reflect motor preparation based on the expectancy and motor execution when making a response that is either or not in line with the expectations. In the comparison of LRP magnitude across task conditions, we found a greater LRP_prep_ when large rewards were more likely, reflecting a greater motor preparation to obtain larger rewards. We also found a greater LRP_exec_ when large rewards were presented unexpectedly, suggesting a greater motor effort placed for executing a correct movement when presented with large rewards. In the analysis of mPFC theta, we found a greater theta power prior to performing an unexpected than expected response, indicating its contribution in response conflict resolution. Collectively, these results demonstrate an optimized motor control to maximize rewards under the dynamic changes of real-life environment.

## Introduction

It is a long-standing belief that rewards drive human behavior (Anderson, 2017; Berridge and Robinson, 1998). In particular, mounting evidence demonstrates that rewards can directly modulate selection and execution of movements (Chen et al., 2018). Modulation by reward is shown for example by reduction in response time and improvement in performance accuracy during oculomotor and reaching movements (Carroll et al., 2019; Galaro et al., 2019; Hickey and van Zoest, 2012; Klein et al., 2012; Summerside et al., 2018; Takikawa et al., 2002), as well as during acquisition and expression of complex motor skills (Chen et al., 2018; Nikooyan and Ahmed, 2015; Wächter et al., 2009). The effect of reward on motor performance is further supported by studies showing reward modulations in neural signals of motor control (Alamia et al., 2019; Bijleveld et al., 2014; Chen et al., 2019; Gluth et al., 2013; Hare et al., 2011; Kapogiannis et al., 2008; Pastor-Bernier and Cisek, 2011; Roesch and Olson, 2003; Sul et al., 2011; Wunderlich et al., 2009) and corticospinal excitability (Galaro et al., 2019; Klein-Flügge and Bestmann, 2012; Klein et al., 2012). These studies demonstrate that presentation of rewards or reward predicting cues increases cortical motor activity which facilitates the movement towards obtaining the reward, also providing support for the idea that the brain motor’s system contributes to value-based decision making (Cisek, 2007, 2006; Cisek and Pastor-Bernier, 2014; Freedman and Assad, 2011; Friston, 2010; Gold and Shadlen, 2007).

Importantly, the dynamic nature of real-life environment imposes several factors that can influence selection and execution of actions. Firstly, the environment is characterized by uncertainty, indicating that we have imperfect knowledge about the outcomes of our actions (Platt and Huettel, 2008). Under these circumstances, we often take actions without knowing whether or not we’ll be rewarded as shown by the example of investing on stocks or pursuing higher education. In such cases, we rely on information about the prospect or expectation of reward delivery to guide our actions (Platt and Huettel, 2008). Another feature of dynamic real-life environment is that it changes constantly and in certain cases, action plans should be flexibly modified to take an alternative action (Krämer et al., 2011; Liebrand et al., 2018; Serrien and Sovijärvi-Spapé, 2013). For instance, take the example of investment decisions. Due to the ever-changing financial market, a once believed ideal stock option can suddenly become devalued. In such scenarios of sudden environment change, your course of action should be adjusted adaptively to maximize rewards in a given state. Adjustment in actions are inevitable especially when considering that we rely on limited information on the prospect or expectations about rewards to guide our behavior and these expectations can sometime be violated (Schultz, 2016).

The current study investigated how the brain’s motor system adapts to the dynamic changes in environment in its pursuit of reward. In particular, using electroencephalography (EEG) we studied how reward magnitude and expectancy influenced the lateralized readiness potential (LRP) (Coles, 1989). Crucially to investigate the effect of sudden environment change on the motor system, we compared the LRPs while participants were performing an expected vs. unexpected action to retrieve a reward. LRPs can be used as an index of activity in the lateralized motor cortex, where the signal is observed as a contralateral negativity that peaks prior to the actual response (Clark et al., n.d.; Mattler et al., 2006; Smulders et al., 2012). It is known to be sensitive to response expectations (Mattler et al., 2006) and a few studies have also shown indication of reward modulation in LRP showing greater negativity for rewarded responses (Pornpattananangkul and Nusslock, 2015; Wang et al., 2019).

Extending onto prior findings, we studied how varying levels of reward magnitude and expectancy modulate LRP patterns. In doing so we leveraged on the high temporal resolution of EEG to uncover the fine-grained temporal cascade of LRP modulation. To this end, we analyzed both the stimulus-locked LRPs reflecting response selection and preparation and the response-locked LRPs reflecting response execution per se (Leuthold et al., 2004, 1996; Osman and Moore, 1993). Additionally, we used a novel analytic approach to further segment the stimulus-locked LRP into an earlier component relevant for processing movement preparation and a later component relevant in movement execution. This new approach was taken based on previous studies showing that an earlier slow wave lateralized activity in stimulus-locked LRP reflected expectations about a response prior to execution (Kemper et al., 2012). By testing the reward modulations in each of the LRP components across time, we determined how rewards shape our motor system across motor hierarchy, while navigating through the dynamic changes in the environment. We hypothesized that reward magnitude and expectancy will modulate LRPs during both the preparation and execution components, suggesting modulation across all levels of motor control.

Apart from LRP, we also investigated the involvement of mediofrontal (mPFC) theta oscillations, known to have significant contributions in both the proactive control of motor plans during action selection (Derosiere et al., 2018; van Driel et al., 2015; van Noordt et al., 2015) and reactive control of readily specified motor commands (Cavanagh et al., 2012; Zavala et al., 2015). We hypothesized that mPFC modulation by task conditions will support its involvement in response conflict resolution (Cavanagh et al., 2010; Töllner et al., 2017; Zavala et al., 2014), further suggesting its role in adaptive control of motor behavior.

## Materials and Methods

### Participants

Thirty-seven right-handed college students (16 males, 19.94 ± 1.16 yrs), without a history of psychiatric and neurological disorders or alcohol/drug dependence, were recruited from the University of Massachusetts Amherst. All study participants signed a written informed consent, approved by the UMass Amherst Institutional Review Board. Participants received course credits or monetary compensation of $15 for participation after completion. In addition to the flat rate of credit/money for participation itself, we also granted a bonus ranging between $1 and $5 based on the reward points they earned throughout the value-based action-selection task.

### Value-based action-selection task

The experiment consisted of two stages – the baseline followed by the task stage. In the baseline stage, participants were presented with a left- or right-oriented grating stimulus (200 msec) (e.g., the 3^rd^ screen in Fig. 1A) and were asked to make a corresponding response within 1000 msec, by pressing either the left key (“A” on the standard 101 keyboard) using their left index finger, or the right key (“L”) using their right index finger. Response times (RT) longer than 1000 msec were considered as incorrect response. Response times from correct responses during the baseline stage were used to calculate the RT categories for determining rewards in the task stage (as in Chen et al., 2019). Four RT categories were determined based on the lognormal distribution of RTs (Category 1: RT < μ – σ; Category 2: μ – σ ≤ RT< μ; Category 3: μ ≤ RT < μ + σ; Category 4: RT ≥ μ + σ; μ and σ refers to the mean and standard deviation of the lognormal distribution). The baseline stage lasted approximately five minutes.

**Figure 1.**
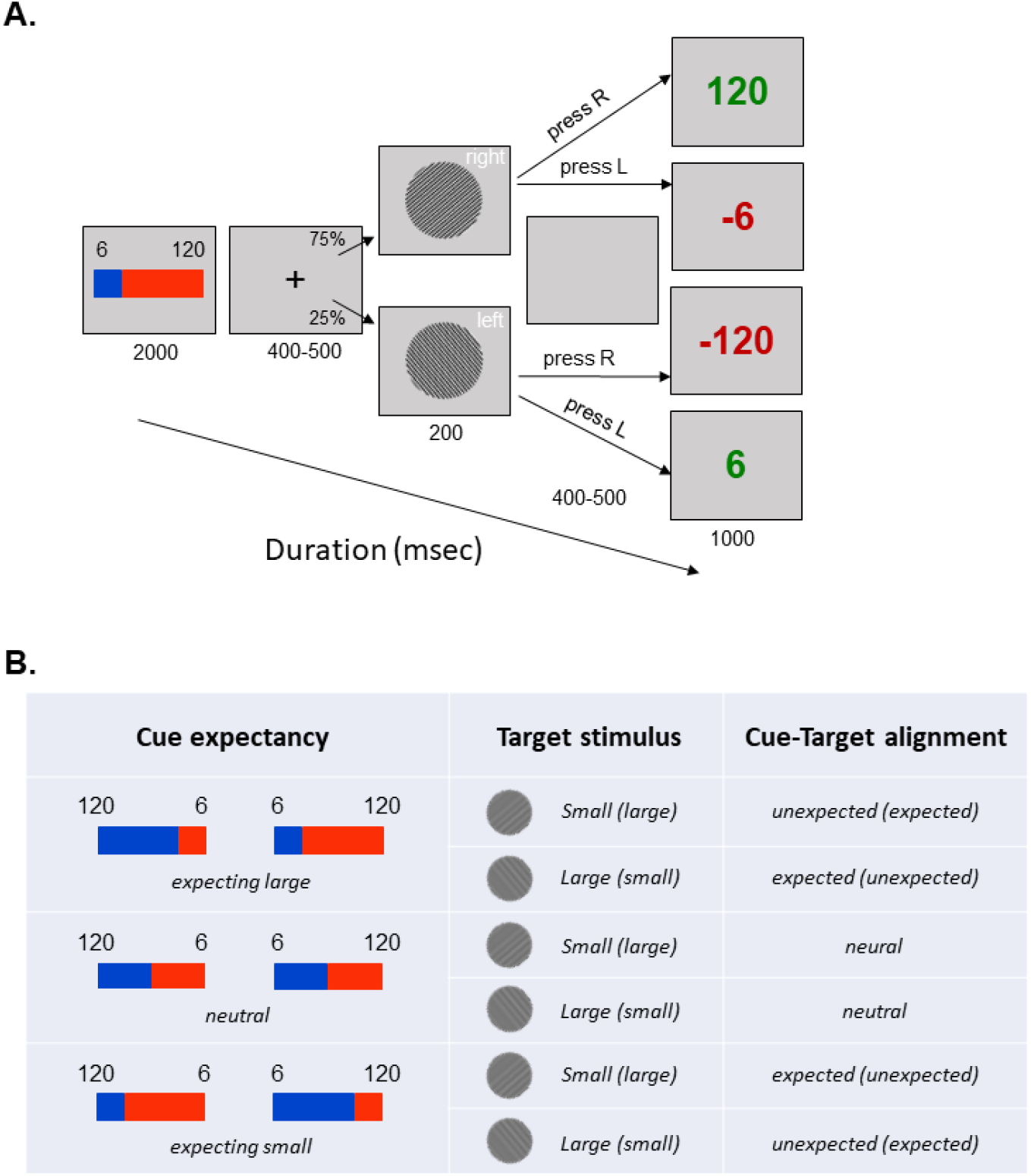
Example trial sequence of the value-based action-selection task (A). Baseline stage of the task only consists of the second to fourth screen on the displayed sequence, during which they make a left- and right-hand response to the left and right grating stimulus. The task stage starts from the presentation of the decision cues and ends with the presentation of the reward feedback. List of task conditions based on the decision cues and target stimulus (B). There were three categories of decision cues based on expectancy (*Cue expectancy*), each followed by the presentation of a grating stimulus that was associated with a small or large reward (*Target stimulus*). Finally, the trials could also be categorized based on the alignment between the cue and the target (*Cue-Target alignment*). For each Cue expectancy condition, there two decision cues to counterbalance the left and right positioning of the small and large reward. The listings in parenthesis is relevant for the decision cue on the right.

In the task stage (Fig. 1A), participants were first presented with a decision cue (2000 msec), which consisted of the reward points (large: 120 or small: 6) associated with the left vs. right grating stimuli and the probability that the left vs. right stimulus will appear (25% vs. 75%, 50% vs. 50% or 75% vs. 25%). Probability assignment was indicated by the area of the red and blue segments in a horizontal bar (Fig. 1B). Combinations of reward and probability assignments resulted in three different expectancies about the target stimulus: 1) expecting large reward stimulus (expecting large), 2) neutral expectation for the two stimuli (neutral) and 3) expecting small reward stimulus (expecting small). To randomize the (left/right) position of relative reward points across trials, we used two sets of decision cues identical in terms of the reward-probability combinations. The two cues only varied in terms of the positioning of reward points. The color of the probability bar (i.e., red and blue) indicating the left and right directions were counterbalanced across subjects to control for color preference.

A variable fixation cross (400-500 msec) was presented after the decision cue, followed by the target left or right grating stimulus determined by the probability assignment. Participants were only allowed to make a response after being presented with the target stimulus. A premature response while still being presented with a decision cue or a fixation cue immediately cancelled the trial. Similar to the baseline stage, a response was required using the corresponding finger within 1000 msec. A correct response was rewarded based on the RT using the pre-defined RT categories determined from the baseline stage. For trials that met the Category 1, 2, 3, and 4, the participants eared 100%, 50%, 25%, 0% of the total number of points assigned to target stimulus, respectively. An incorrect response resulted in loss of total points assigned to the side of the key press (i.e. resulted in -120 or -6). For example, in Figure 1A, if a participant made a correct right-hand response to a right-oriented stimulus, a proportion of 120 points determined by the speed of response was granted. If the participant made an incorrect right-hand response to a left-oriented stimulus, 120 points assigned to the right-hand response were lost (i.e., -120 points). Earned and lost points were displayed in green and red respectively at the end of each trial (1000 msec), followed by a variable inter-trial interval (400-500 msec). Participants completed total of 480 trials across 10 blocks. Within a block, each of the six decision cues were chunked into a semi-block with 4-12 trials which consisted of the same cue. Inclusion of semi-blocks was to make the processing of the decision cues easier by repeating the same cue several times. The order of semi-blocks was randomized within each block. Each of the six cues were presented in 80 trials throughout the experiment. After each block, participants were shown the accumulated amount of points they have earned up until then. The task stage lasted approximately 50 min.

### EEG recording and analysis

The electroencephalogram (EEG) was continuously recorded using 64 scalp electrodes embedded in an extended coverage, triangulated equidistant cap (M10, EasyCap, GmbH) using a low-pass filter of 100 Hz at a sampling rate of 1000 Hz (actiCHamp, Brain Products, GmbH). The electro-oculogram (EOG) was monitored with electrodes below the left eye and just lateral to the left and right canthi. In rare cases, channel impedances up to 25 kΩ were tolerated, but in most cases, they were kept below 15 kΩ. The EEG was amplified with a BrainAmp system (Brain Products GmbH, Gilching, Germany). All channels were referenced to the vertex (Cz) during recording.

Offline EEG data was exported to Matlab using the EEGLAB software package (Delorme and Makeig, 2004), and custom scripts. The data were re-referenced to the average of mastoid channels and were high-pass filtered at 0.1 Hz. We created two epochs– one time-locked to the target stimulus (−250 to 1000 msec) and one time-locked to the response (−1000 to 500 msec).

For each participant, we implemented a procedure for artifact removal based on an independent component analysis (ICA) approach (Delorme et al., 2012; Makeig et al., 2004; Onton and Makeig, 2006) that has been known to greatly diminish the contribution of ocular/biophysical artifacts. Single trials were also visually inspected to exclude epochs with excessively noisy EEG or muscle artifacts. On average, 96.98% of the stimulus-locked epochs and 95.88% of the response-locked epochs were included in the final analysis.

For the analysis of dynamics in neural oscillations, oscillatory power of was calculated by means of Fast Fourier Transformation (FFT) as implemented in the new*timef* function in EEGLAB. This procedure implements the calculation of changes in oscillatory power by means of a moving window (steps = 10 msec). The stimulus locked, and response locked data (both oscillatory responses and ERPs) were subsequently binned (excluding trials during which an incorrect response was given) and averaged within each task condition.

To determine the fine-grained temporal cascade of LRPs, we identified the LRP in both the stimulus-locked and response-locked data. We focused on the following cortical motor regions of interest (ROI) on each side of the hemisphere: left motor (FC3, FC5, C3, C5), right motor (FC4, FC6, C4, C6) (Deiber et al., 2012; Gregory et al., 2016; López-Larraz et al., 2015; Picazio et al., 2014). Lateralized motor signals were identified by contrasting the ERPs of the ipsilateral from contralateral motor ROI relative to the hand that was used to respond (Fig. 2). Furthermore, to study the engagement of mediofrontal theta oscillations reflecting cognitive control, we studied the oscillatory signal from the mediofrontal ROI (FCz, Fz) (Frank et al., 2015; Mas-Herrero et al., 2015). Analysis if oscillatory power was conducted using cluster-based permutation testing (Maris and Oostenveld, 2007) across frequency and time points within a specified time range (−400-0 msec from response onset).

**Figure 2.**
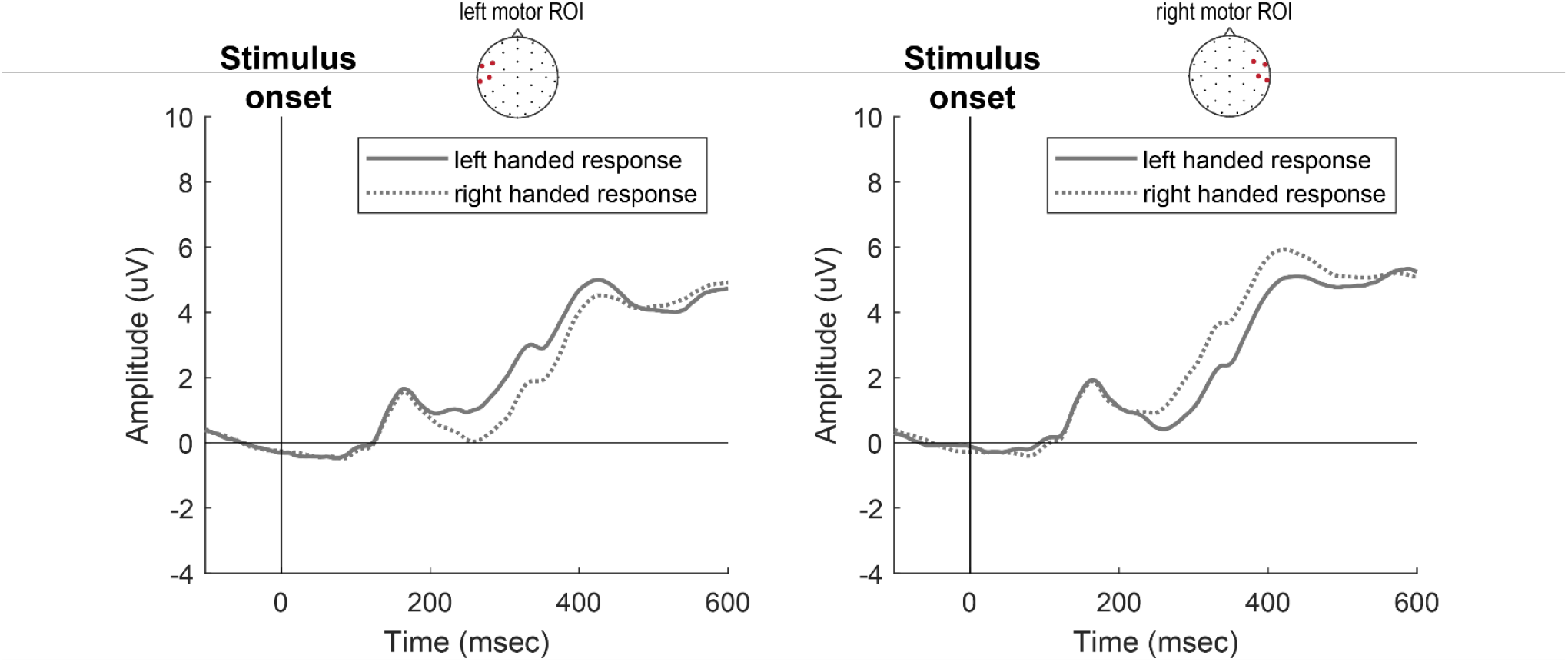
Grand average ERP traces from the left and right motor ROI

### Statistical analysis

Analyses of the behavioral and EEG data were focused on determining the effect of the three decision cues reflecting decision scenarios that varied in terms of reward magnitude and expectancy: “expecting large”, “neutral”, “expecting small” (Fig. 1B). Following the decision cue, an actual target of either a “large” of “small” reward stimulus appeared to which a response was made. Depending on the alignment between the cue expectancy and target stimulus, each outcome can be considered as “expected” (e.g., small reward stimulus after expecting small cue), “unexpected” (e.g., large reward stimulus after expecting small cue) or “neutral” (e.g., either small or reward stimulus after a neutral cue). For comparisons across all of the outlined task conditions, we used a repeated measures ANOVA model (with type 3 sums of squares) (Herr, 1986) including three main effects: Cue expectancy, Target stimulus and Cue-Target alignment (see Fig. 1B for the different level of each factor). In addition, we added a term for the interaction between Cue expectancy and Target stimulus and between Target stimulus and Cue-Target alignment, to specifically focus on how the effect of stimulus reward magnitude was modulated by prior expectation based on the cues as well as whether or not these expectations were met. ANOVAs are reported with Greenhouse-Geisser corrected p-values (Greenhouse and Geisser, 1959) if sphericity assumptions were violated when tested using Mauchly’s W test (Mauchly, 1940). As mentioned, there were two sets of decision cues that varied only in terms of right and left positioning of reward points. The two sets were combined for all the behavioral (RTs and accuracy) and neural measures prior to the analysis across conditions. For simplicity we only report significant F test in all our results, unless mentioned otherwise.

## Results

### Behavioral results

Analysis of RT (Fig. 3A) showed a main effect of Target stimulus suggesting faster response to a large (M = 342, SE = 5.9) as compared to a small (M = 354, SE = 5.5) reward stimulus (*F*(1,36) = 36.19, *p* < .001). We also found a main effect of Cue-Target alignment (F(2,72) = 92.81, p_gg_ < 0.0001) suggesting faster responses to expected (M = 332, SE = 6.1) as compared to unexpected (M = 364, SE = 5.8) condition and the neutral (M = 349, SE = 5.4) condition in between (all post hoc comparisons p < .001). Finally, we observed a significant Cue expectancy by Target stimulus interaction (*F*(2,72) = 101.97, *p*_*gg*_ < .001). Post-hoc analysis showed that after the large reward cue, as well as following the neutral cue, response was significantly faster to the large than small reward stimulus (expecting large cue: M_large_ = 359, SE = 6.4, M_small_ = 368, SE = 5.8; neutral cue: M_large_ = 343, SE = 5.8, M_small_ = 354, SE = 5.2, both *p*s < .001). After the small cue, responses were faster to small (M = 339, SE = 6.0) than large (M = 359, SE: 6.2) reward stimulus (*p* < .001).

**Figure 3.**
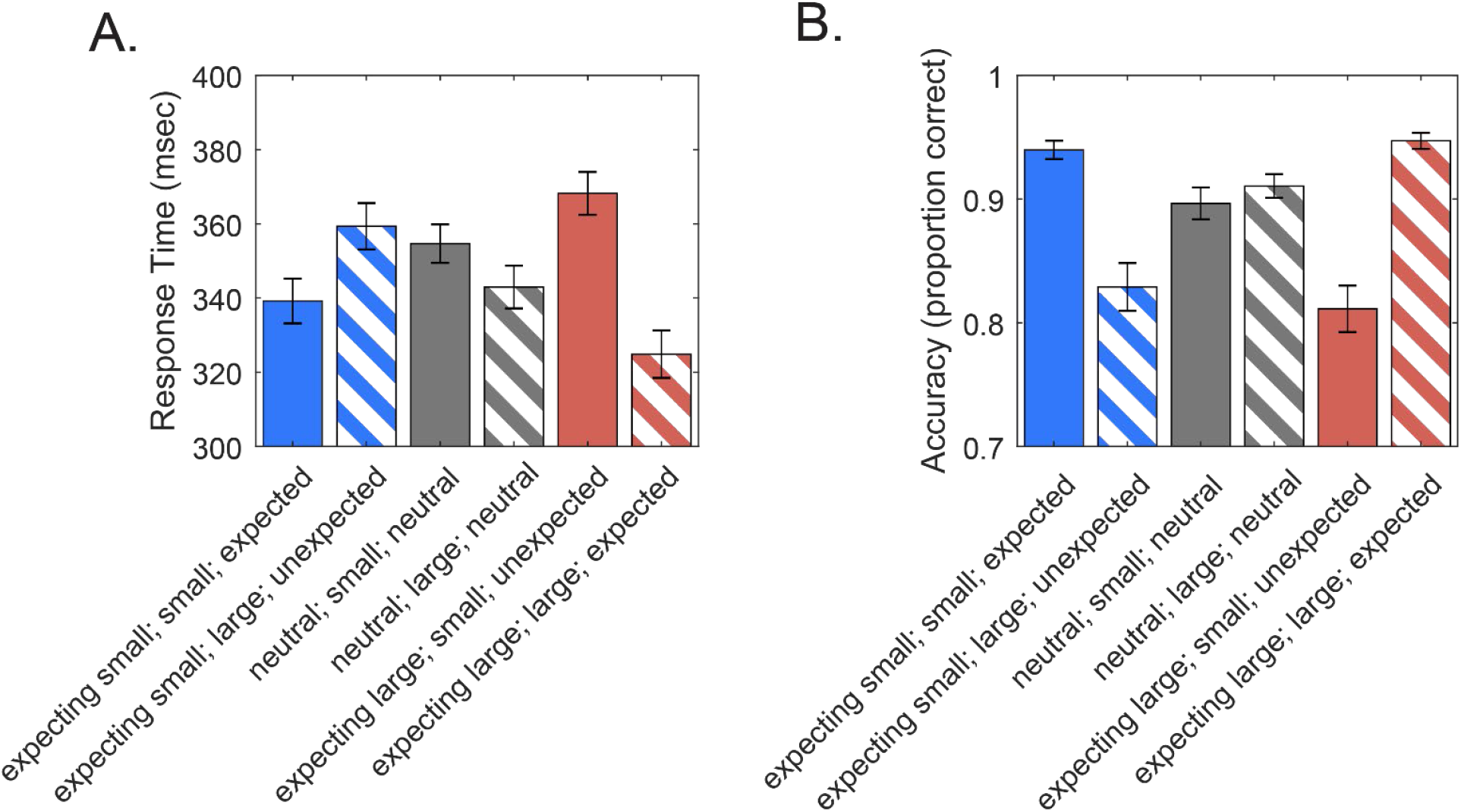
Response time (A) and accuracy (B) in each task condition. The labeling of task conditions are based on the combination across different levels of the factors, Cue expectancy, Target stimulus and Cue-Target alignment in serial order [Cue expectancy; Target stimulus; Cue-Target alignment]. Solid and grated bars denote small and large reward target stimulus conditions; blue, gray and red denote expecting small, neutral and expecting large cue expectancy conditions, respectively.

A similar pattern was found in the analysis of accuracy (Fig. 3B). Here we found a significant main effect of Cue expectancy (*F*(2,72) = 8.83, *p*_*gg*_ < .001), suggesting higher accuracy in neutral as compared to the expecting small (p = 0.013) and expecting large reward cue (p = 0.0002) (M_expecting small_ = .88, SD = .07, M_neutral_ = .90, SE = .01, M_expecting large_ = .88, SE = .01). We also found a main effect of Cue-Target alignment (F(2,72) = 64.7, p_gg_ < .001) which suggested higher accuracy for expected (M = .94, SE = .01) compared to unexpected (M = .82, SE = .02) condition and the neutral (M = 0.90, SE = .01) condition in between (all post hoc comparisons p < .001). Finally, we found Cue expectancy by Target stimulus interaction (*F*(2,72) = 61.51, *p*_*gg*_ < .001) (Fig. 3B). Post-hoc analysis showed a significantly higher accuracy for large (M = .95, SE = .01) than small reward stimulus (M = .81, SE = .02) (*p* < .001) after expecting large cue. After expecting a small cue, accuracy was significantly higher for small (M = .94, SE = .01) than large reward stimulus (M = .83, SE = .02) (*p* < .001). No significant difference was found in the neutral condition (*p* = .154).

### Stimulus-locked LRP (_sl_LRP)

The stimulus-locked data showed an _sl_LRP (Fig. 5A and B) starting at ∼175msec, and peaking at ∼320msec. No significant condition effects were found on _sl_LRP peak latency (all Fs < 1.56, p > 0.21). The peak amplitude (averaged across 315 ± 20msec) of the _sl_LRP was slightly larger (more negative) for unexpected as compared to expected condition (main effect of Cue-Target alignment: F(2,72) = 3.57, p_gg_ = 0.034; all other F < 2.2, p > 0.12). Furthermore, there seemed to be some reward magnitude and expectancy modulations preceding 175msec. For instance, following an expecting large cue, we observed a contra-lateralized negative variation when the target stimulus also gave a large reward. However, with a small reward stimulus after the same “expecting large” cue, we observed a positive variation followed by a sharp change from positive to negative polarity. This reflects an increased motivation to use the other hand (assigned with the large reward), before a sudden change in movement plan for execution of correct response. This suggest that _sl_LRP waveform reflects at least two separate motor processes: 1) continuation of motor preparation based on the preceding decision cue and 2) motor execution after the identification of the actual target stimulus.

**Figure 4.**
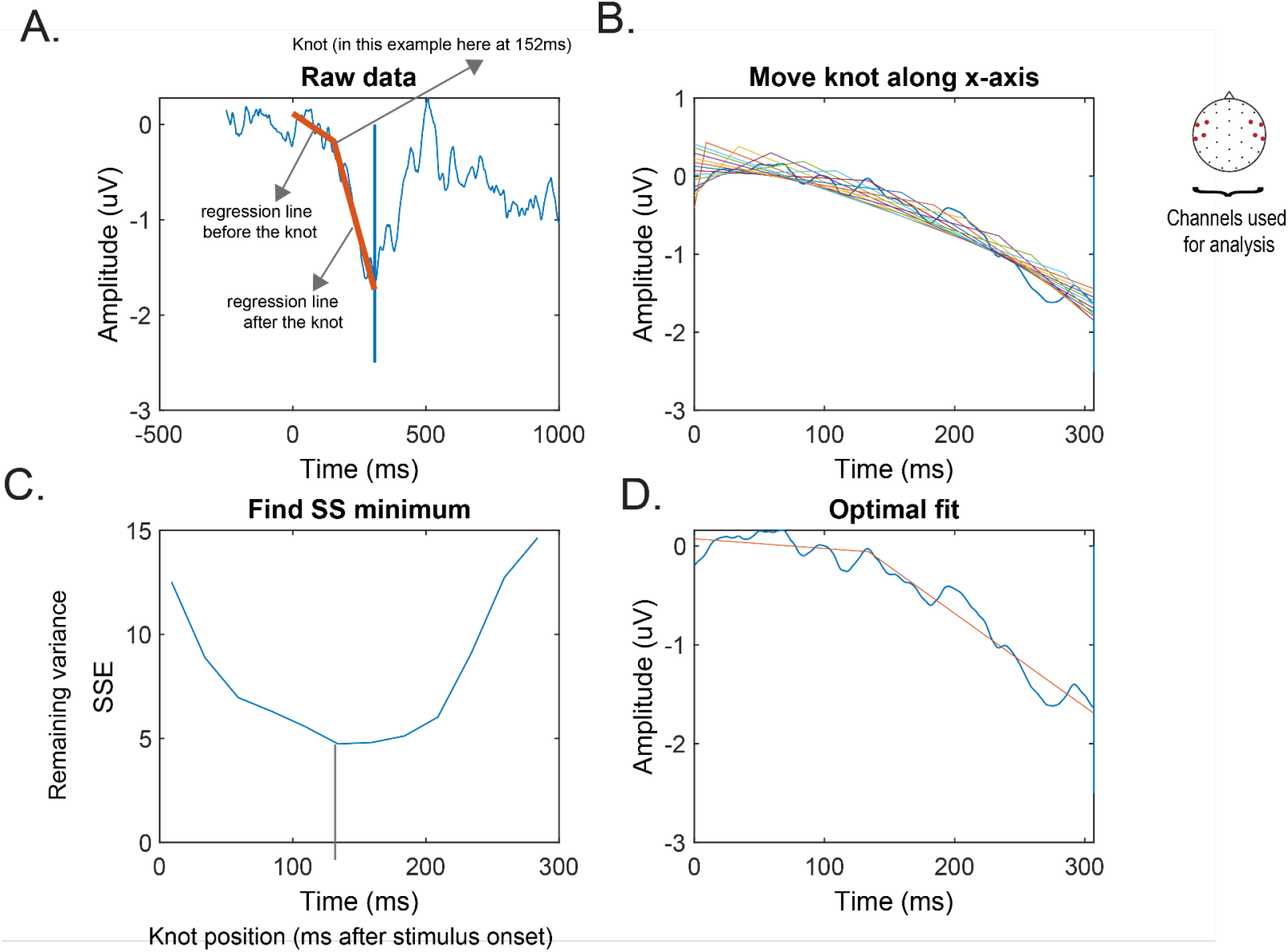
Procedures for segmenting the stimulus locked LRP (Time 0 = stimulus onset) into preparation and execution components using piecewise regression. Two linear lines were fit to the raw stimulus-locked LRP (A). The two lines met at a knot, where the slope of the two lines together explained most variance(B-D).

**Figure 5.**
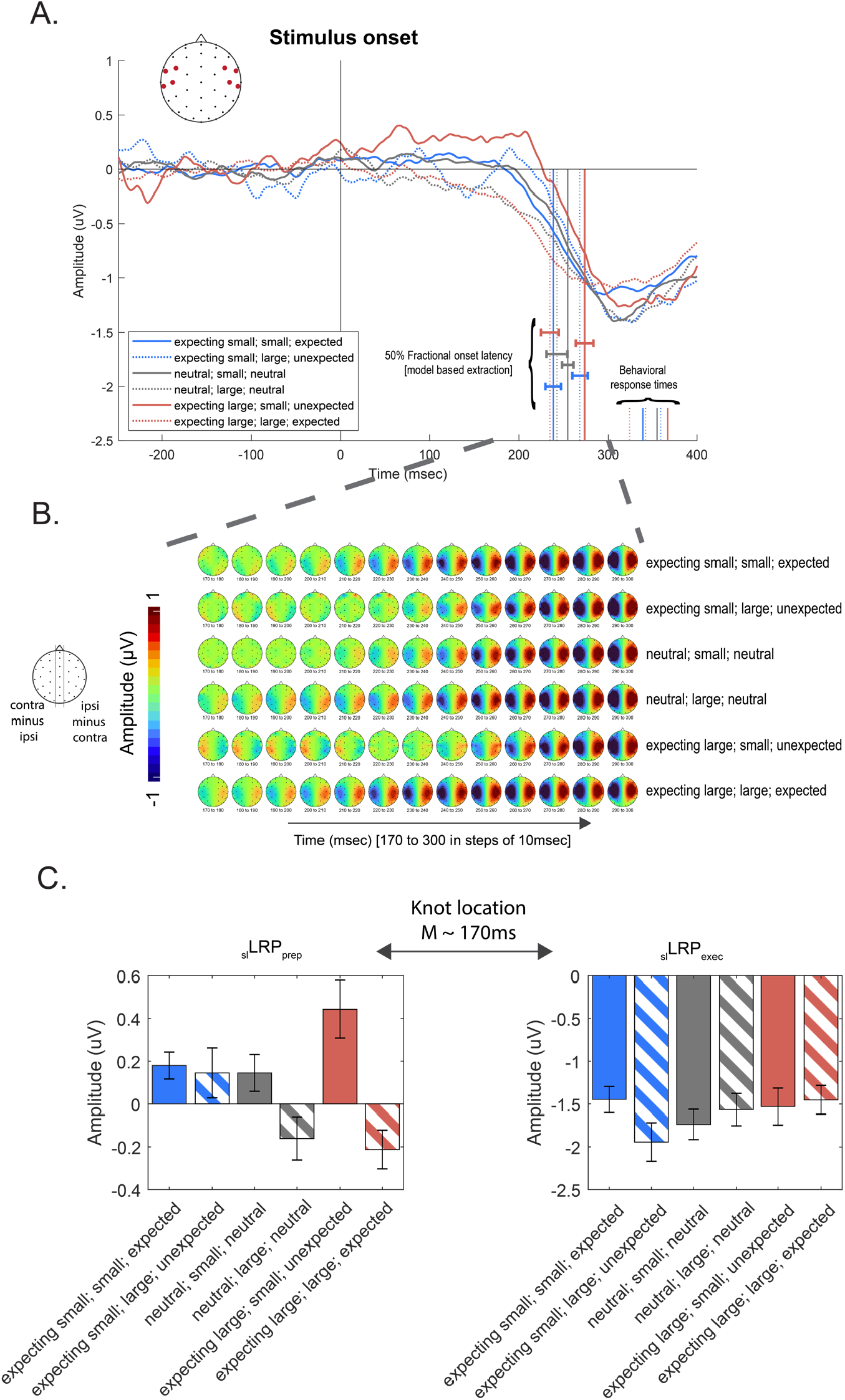
Evoked lateralized motor responses time-locked to the stimulus (_sl_LRP). LRP was extracted from the motor ROI by subtracting the ipsilateral from the contralateral hemisphere relative to response (A). The LRP onset (as determined by the 50% fractional peak latency) is depicted by the set of vertical lines between 200 and 300 msec. The vertical lines between 300 and 400 msec depict response time in each condition. The topographical plot of LRP across time (B). The different rows depict each task condition. Comparison of LRP amplitude across task conditions in the preparation (_sl_LRP_prep_) and execution (_sl_LRP_exec_) component (C)

To model these two separate motor processes in the _sl_LRP waveform we used a piecewise regression method (Rashid et al., 2019). Two linear regression lines were fitted through the stimulus locked LRP averaged within each condition in each individual (Fig. 4A). The starting point of the first line was set at stimulus onset (0 msec – the intercept) and the ending point of the second line was set at the LRP peak latency, determined based on the average across all conditions within an individual (since peak latency was not significantly influenced by condition, collapsing within individual improved our power to detect the LRP peak latency; between 0 and 400msec). The two regression lines met at a knot. The location of the knot was optimized by moving the knot position along the x-axis (i.e., across time) and identifying the time point that resulted in the least amount of unexplained variance by the two regression lines (Fig. 4 B-D). This approach to independently model motor preparation and execution allows for efficient isolation of the slow wave activity preceding the actual _sl_LRP dip, which is known to be influenced by expectations related to preparing a specific motor response (Kemper et al., 2012; Mattler et al., 2006).

After decomposing the stimulus-locked LRP waveform into preparation (_sl_LRP_prep_) and execution (_sl_LRP_exec_) related components, we separately investigated the task condition effects on each component. We used the slope of each of the two regression lines as a proxy for amplitudes of _sl_LRP_prep_ and _sl_LRP_exec_. The _sl_LRP_prep_ showed a significant main effect of Target stimulus (F(1,36) = 19.99, p < 0.001); the _sl_LRP_prep_ was more negative for large (M = -0.08, SE = 0.056) than small (M = 0.26, SE = 0.051) reward stimuli (Fig. 5C), suggesting a preparational bias towards the large reward regardless of expectancy. We also found a significant main effect of Cue-Target alignment (F(2,72) = 5.47, p_gg_ = 0.007), with the _sl_LRP_prep_ being more positive (i.e. reflecting preparation effects of the other hand) for unexpected (M = 0.29, SE = 0.087) compared to expected (M = -0.016, SE = 0.0533) condition and the neutral condition in between (M = -0.0077, SE = 0.0736) (expected vs. unexpected: p = 0.0099; neutral vs unexpected: p = 0.042). The pattern is in line with our task manipulations, demonstrating response preparation towards the other side when the cue and the target was not aligned. Finally, we observed a significant Cue expectancy by Target stimulus interaction (F(2,72) = 5.04, p_gg_ = .009). Post-hoc analysis showed that the _sl_LRP_prep_ was significantly more negative for large than small reward stimulus after the expecting large cue (p < .0001, M_large_ = -.21, SE = .09, M_small_ = .44, SE = .13) and the neutral cue (p = .012, M_large_ = -.16, SE = .10, M_small_ = 0.15, SE = 0.09). No difference in _sl_LRP_prep_ amplitude was observed following the expecting small cue. These results indicate a stronger preparational bias towards the large reward side when expecting large rewards and even in neutral expectancy, while no such bias is detected when expecting small rewards.

For the _sl_LRP_exec_ we found a significant main effect of Cue-Target Alignment (F(2,72) = 3.4, p_gg_ = 0.039), with the _sl_LRP_exec_ being more negative for unexpected (M = -1.73, SE = 0.18) compared to expected (M = -1.45, SE = 0.15) condition and the neutral condition in between (M = -1.65, SE = 0.16) (unexpected vs expected: p = .043; no other pairwise comparisons significant, all other ps > 0.16 (Fig. 5C). Furthermore, we observed a significant Cue expectancy by Target stimulus interaction (F(2,72) = 4.82, p_gg_ = .011). After expecting small cue, the _sl_LRP_exec_ was larger (more negative) for the large (M = -1.94, SE = 0.23) as compared to the small (M = -1.45, SE = 0.15) reward stimulus (p = 0.006). No such difference was found in expecting large or neutral cue (p > 0.3). The Target stimulus by Cue-Target alignment interaction was marginally significant (F(2,72)=2.4, p_gg_ = 0.08). Here, an exploratory follow up tests revealed a marginal difference between unexpected small and large reward stimuli (p = 0.096). These results suggest a greater motor effort being placed to execute a correct response specifically when a large reward stimulus appeared unexpectedly.

After investigating the amplitudes of _sl_LRP components, we also tested for the difference in the timing of the knot which is at the end of _sl_LRP_prep_ and the start of _sl_LRP_exec_ (see Fig. 4 for approaches identifying knot timing). This time point will reflect relative timing of preparation vs. execution process. A significant effect of Cue-Target alignment (F(2,72) = 3.59, p_gg_ = 0.0326) suggested that the _sl_LRP knot location was earlier for expected (M = 171, SE = 8.42) (p = .056), and neutral (M = 174, SE = 7.36) (p = .062) as compared to unexpected (M =197, SE = 8.51) condition. Furthermore we observed a Cue expectancy by Target stimulus interaction (F(2,72) = 5.89, p = .0048). When expecting small, the knot location was later for large (M = 210, SE = 11) than small reward (M = 160, SE = 10.70) stimuli (p = .0012). Such difference was not found in expecting large and neutral cue expectancy conditions (all ps > 0.13). We also found a significant Target stimulus by Cue-Target alignment interaction (F(2,72) = 3.69, p = .0298). When Cue-Target alignment was “unexpected”, the difference between the large (M = 210, SE = 11) and small reward (M = 185, SE = 11) stimuli was marginally significant (p = 0.07), with the knot location happening later for large reward stimulus. No such difference was found in neutral and expected conditions (all ps > 0.13).

Next, we used the combination of the two slopes to model when the 50% fractional LRP onset latency (i.e. when the _sl_LRP amplitude was equal to 50% of its peak amplitude (Kiesel et al., 2008; Mordkoff and Gianaros, 2000). 50% peak amplitude latency has been previously used to estimate the start of the LRP. The vertical color-coded vertical lines between 300 and 400 msec in Fig. 5A depict the extracted onset latency. The pattern of the stimulus locked LRP onset matched the pattern as observed in the actual behavioral responses (smaller color-coded vertical lines labeled as “behavioral response times” between 300-400 msec in Fig. 5A). Statistical analysis revealed a significant effect of Cue-Target alignment (F(2,72) = 10.31, p_gg_ = .0001). Here, the LRP onset was much earlier for expected (M = 237, SE = 7.2, SD = 43) than unexpected (M = 271, SE = 7.46) stimuli (p < 0.001), with the onset following neutral cue in between (M = 249, SE = 7.36). We also found a Cue expectancy by Target stimulus interaction (F(2,72) = 8.75, p_gg_ = 0.00048). When expecting large, the LRP onset was earlier for large (M = 235, SE = 10) than small reward (M = 274, SE = 10) stimulus (p = .005). However, when expecting small, the onset was later for large (M = 269, SE = 8) than small reward (M = 239, SE = 9) stimuli (p = .005). Such difference was not found in neutral cue expectancy conditions (p> 0.32).

### Response-locked LRP (_rl_LRP)

In the response-locked data, we observed the <u>_rl_LRP</u> peak around 40 msec prior to response (Fig. 6). The <u>_rl_LRP</u> amplitude was subsequently extracted between -60 and -20 msec prior to the given response (peak latency ± 20msec) (Luck, 2014). We found a significant Cue expectancy by Target stimulus interaction effect (F(2,72) = 5.294, p_gg_ = 0.014). After expecting small cue, <u>_rl_LRP</u> was more negative for large (M = -2.31; SE = 0.28) than small (M = -1.81; SE = 0.19) reward stimulus (p = 0.012). No such effect was found after expecting large or neutral cues (all ps > 0.23). These results are parallel to the execution component of stimulus-locked LRP (_sl_LRP_exec_), suggesting that a greater motor effort is placed when a large reward stimulus appeared unexpectedly.

**Figure 6.**
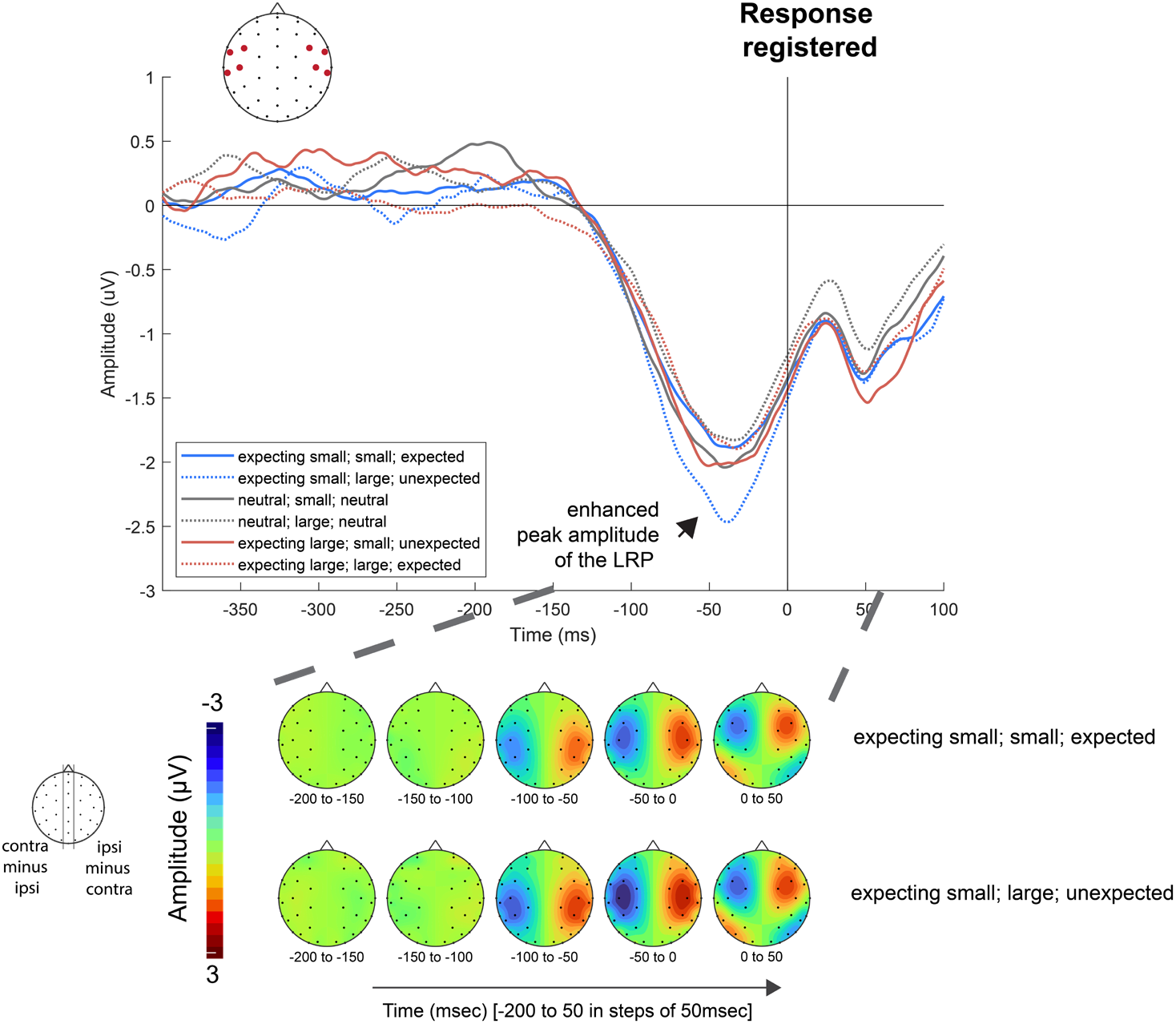
Evoked lateralized motor responses time-locked to response (_rl_LRP). Similar to _sl_LRP, we extracted LRP by subtracting the signal in ipsilateral from the contralateral motor ROI relative to response. The LRP amplitude preceding the response (between -60 and -20 msec) when a large reward stimulus appeared unexpectedly.

### Neural oscillations

After studying the LRP, we investigated the involvement of mediofrontal theta oscillations to test for the prefrontal cognitive control mechanism during motor selection and execution (Cavanagh et al., 2012; Derosiere et al., 2018; van Driel et al., 2015; van Noordt et al., 2015; Zavala et al., 2015). Starting around 200msec prior to response (Fig. 7A), oscillatory theta power (4-7Hz) was higher for unexpected compared to expected condition (cluster-based correction p < 0.001) indicative of increased cognitive control due to a response conflict resulting from the discrepancy between the cue and the target (Fig. 7B). Confirming this, a repeated measures ANOVA on mean theta power from -200 until response showed a significant effect of Cue-Target alignment (F(2,72) = 20.3, p_gg_ < .0001), with higher theta power for unexpected (M = 0.872, SE = 0.019) compared to both neutral (M = 0.846, SE = 0.019) and expected (M = 0.841, SE = 0.019) condition (both ps < .001; difference between expected and neutral p > 0.34). There was also a significant interaction between Cue expectancy and Target stimulus (F(2,72) = 22.1, p_gg_ < .0001). When expecting large, theta power was lower for large (M = 0.839, SE = 0.019) than small reward (M = 0.873, SE = 0.019) stimulus (p < 0.001). However, when expecting small, theta power was higher for large (M = 0.87, SE = 0.019) than small reward (M = 0.843, SE = 0.019) stimuli (p < 0.001). Such difference was not found in neutral cue expectancy conditions (p > 0.24).

**Figure 7.**
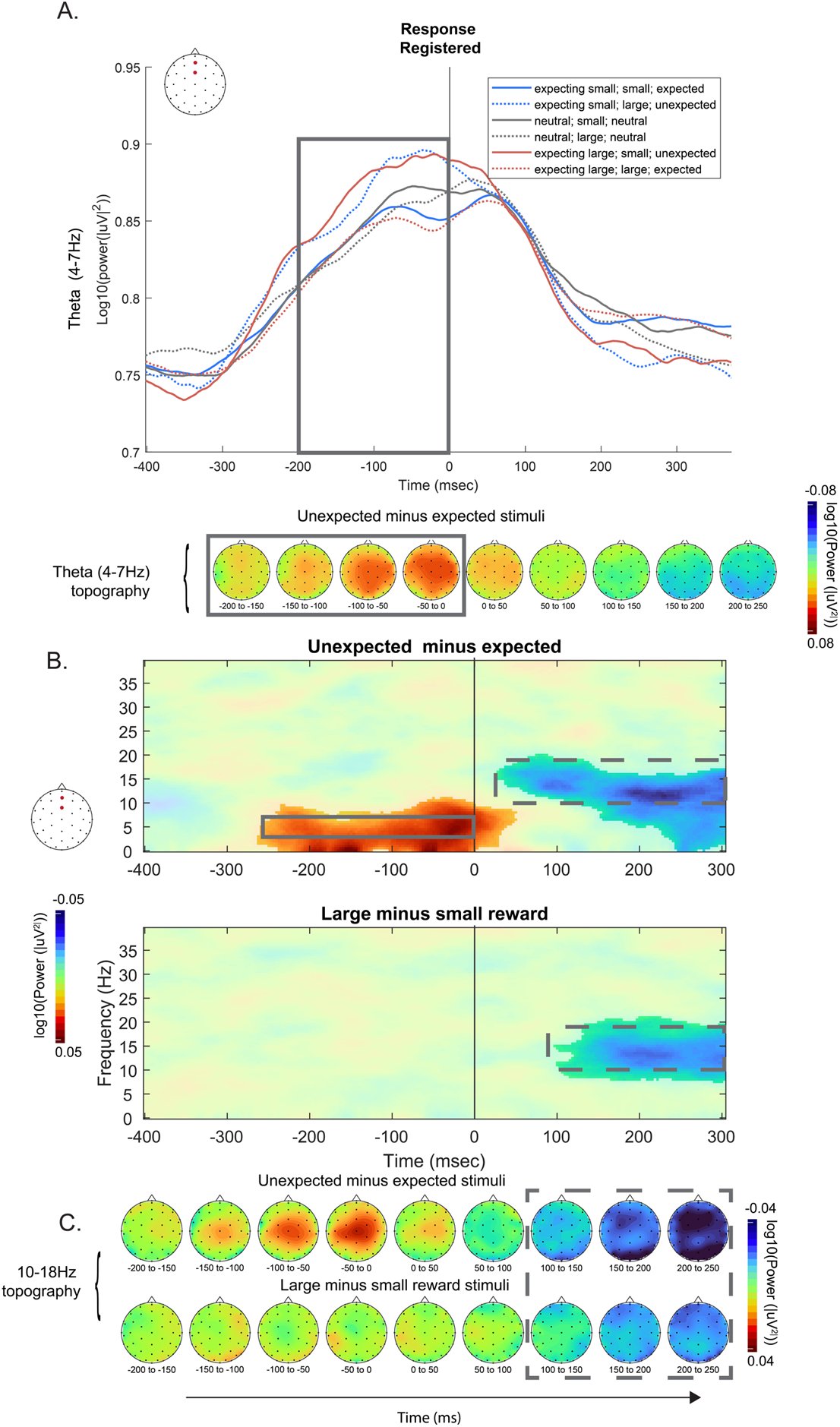
Response locked oscillatory responses. Theta power (4-7Hz) extracted from the frontocentral ROI (A). Prior to the response, theta power was higher for unexpected compared to expected stimuli (solid grey rectangle). Cluster based permutation testing reveal significant differences for the effect of expectancy in the 4-7Hz frequency range, indicative of theta modulation due to stimulus response conflict (B). After the response there was significant decrease in high alpha / low beta (10-18Hz range) for both the effect of reward and expectancy (dashed grey rectangles). Topographical distribution of high Alpha/ low Beta power for the effect of expectancy (top), and reward (large minus small) (bottom) (C).

Additionally, we observed decreased power in the high alpha/low beta range (10-18 Hz) for unexpected compared to expected condition (cluster based p < 0.001), starting around 50msec after response and for large compared to small reward stimulus (cluster based p < 0.001), starting around 100 after response (Fig. 7B and C). An exploratory repeated measures ANOVA on mean power from 50 until 250msec post response revealed a significant main effect of Target stimulus (F(1,36) = 20.11, p < 0.001) with higher alpha/beta power for small (M = 0.483, SE = 0.02) compared to large reward (M = 0.501, SE = 0.02). Furthermore there was also a main effect of cue-target alignment (F(2,72) = 17.2, p_gg_ < 0.001) with high alpha / low beta for unexpected (M = 0.477, SE = 0.02) compared to both neutral (M = 0.497, SE = 0.02) and expected (M = 0.50, SE = 0.02) condition (both ps < .001; difference between expected and neutral p > 0.15). There was also a significant interaction between Cue expectancy and Target stimulus (F(2,72) = 15.1, p_gg_ < 0.001). When expecting large, alpha power was higher for large (M = 0.493, SE = 0.02) than small reward (M = 0.483, SE = 0.02) stimulus (p < 0.001). However, when expecting small, alpha power was lower for large (M = 0.471, SE = 0.02) than small reward (M = 0.512, SE = 0.022) stimuli (p < 0.001). Such difference was not found in neutral cue expectancy conditions (p > 0.15).

## Discussion

Rewards have strong modulatory effect on human behavior (Berridge, 2012; Berridge and Robinson, 1998). Much research to date shows evidence of direct reward modulation on movement selection and execution (Chen et al., 2018). The current study investigated the effect of reward on the brain’s motor system by probing the ERP component, Lateralized Readiness Potential (LRP). In our study, we focused on the fact that in real life, pursuit and retrieval of rewards happen in an environment characterized by uncertainty and dynamic changes. To model the behavior under such circumstances, we compared neural processes related to response preparation and execution across task conditions where participants were presented with decision cues that varied in term of the reward magnitude and expectancy. After the decision cue, participants were presented with a target stimulus that were either or not in line with the expectations based on the probabilistic nature of the cue.

Our behavioral data confirmed that our task conditions captured the effect of reward magnitude and expectancy as well as how much the expectations from the cue were aligned with the target stimulus. Participants responded faster to a large reward stimulus and their responses were more accurate when the stimulus came after a non-biasing (neutral) cue. Responses were both faster and more accurate when the expectations were aligned with the target. The observed modulation of motor behavior by reward magnitude and expectancy can happen at multiple levels of motor control from movement selection, planning to execution. To explore the motor system across these levels, we studied the effect of task conditions across fine-grained temporal windows of LRP, leveraging on the high temporal resolution of the ERP signal time locked to both the stimulus onset and response registration.

Previous works have suggested that while the stimulus-locked LRP reflects response selection and preparation, the response-locked LRP reflects specifically response execution (Leuthold et al., 2004, 1996; Osman and Moore, 1993). These studies also suggesting functional differences that are in line with the levels of motor control or motor hierarchy (Grafton and Hamilton, 2007; Haruno et al., 2003; Uithol et al., 2012).

In looking at the stimulus-locked LRP, we found noticeable task condition effects on an earlier slow wave component. Previous studies have reported that lateralized slow wave motor activity in stimulus-locked signal are influenced by expectations related to preparing a motor response (Kemper et al., 2012; Mattler et al., 2006). To tease apart this slow wave activity from the rest of the stimulus-locked LRP, we applied a piecewise regression approach and identified a knot that reflects the end of preparation and start execution. We analyzed the amplitudes of both the preparation and execution components as well as the knot timing.

We found a differential pattern of effects on the preparatory and execution components of lateralized motor activity in terms of the cue and target stimulus alignment. While in the preparation component LRP amplitude were more positive in the unexpected compared to neutral and expected condition, the pattern was reversed in the execution component with greater negativity observed in the unexpected followed by neutral and expected condition. This reflects preparation towards the other hand followed by a greater effort in making a response when the expectations about the target were not met. This contrasting pattern confirmed that the preparation and execution components have different functional implications, with the former reflecting a continuation of motor preparation based on the decision cues and the latter reflecting the motor efforts after identifying the target stimulus. Additionally, the preparation component showed a stronger preparation towards the large reward side, especially when expecting a large reward stimulus as well as in neutral expectancy condition. No such bias towards one side was shown when expecting a small reward stimulus, suggesting no preparational bias when a small reward was highly likely. These results collectively demonstrate a strong preference for high reward at the stage of movement preparation.

In the execution of component of LRP, we also found an interaction between decision cue and stimulus reward levels. Significant differences across reward levels were only found after the expecting small cue. LRP amplitude was larger when a large reward stimulus followed the expecting small cue, which is in line with the results showing greater LRP (i.e., more negativity) when expectations were not aligned with the target as described above. Interestingly, such difference across stimulus reward levels was not significant when following the expecting large cue. In other words, a significantly greater motor effort was placed to correct for the movement plan only when confronting a large reward after expecting small but not when confronting a small reward after expecting large. This is also supported by the Target stimulus by Cue-Target alignment interaction, showing larger LRP in large than small reward, particularly when the cue and target was not aligned. A similar pattern was observed with response-locked LRP amplitude. Together these results demonstrate significant modulation of motor efforts by reward magnitude and expectancy. The results from the preparation and execution components of stimulus-locked LRP extend the larger literature on LRP, by demonstrating how the discrete shape and features of LRP depicts modulation of motor behavior by for example, response complexity, expectancy and advance preparation (Hackley and Miller, 1995; Hsieh and Yu, 2003; Leuthold, 2003; Masaki et al., 2004; Mattler et al., 2006; Meckler et al., 2010; Müller-Gethmann et al., 2003).

In addition to examining the LRP amplitudes, we compared the timing of different LRP indices. First, we looked at the timing of the knot in stimulus-lock LRP, which reflects the transition from preparation to execution. Overall, we found a later knot timing when cue and target were not aligned and when confronting a large reward stimulus that was unexpected, which is in line with the LRP magnitude results. The later knot timing in these conditions may imply greater amount of mental processing required before transitioning from preparation to execution of response. These mental processes may involve cognitive control related with switching over from previous motor plans to an alternate plan which may also be influenced by reward magnitude (Kenner et al., 2010; Krämer et al., 2011; Liebrand et al., 2018; Rangel-Gomez et al., 2015; Serrien and Sovijärvi-Spapé, 2013). We also analyzed the LRP onset latency and, as opposed to the knot timing, found an almost identical pattern to response time. This is in line with the previous discussions about the implications of LRP latency on the timing of motor preparation and execution (Mordkoff and Gianaros, 2000; Schwarzenau et al., 1998; Smulders et al., 2012). Our results suggest that the LRP latency indeed can accurately estimate the timing of behavioral responses. Furthermore, these findings provide additional support for knot timing as a distinctive measure reflecting mental process that happens prior to response execution.

Apart of the analysis of LRP, analysis of neural oscillations gave us further insights about how the prefrontal cognitive control mechanisms contribute to motor selection and execution. The mediofrontal theta oscillations in particular, was significantly increased when confronting an unexpected than expected cue-target alignment. Importantly this increased theta oscillations appeared before response—around -250 msec from when response was registered—and disappeared shortly after the response was registered. This indicates that the role of mediofrontal theta is involved when response conflict is present to optimize the forthcoming response, replicating prior studies (Gbadeyan et al., 2016; Miller and Cohen, 2001; Stokes et al., 2013). The fact that the mediofrontal theta and LRP modulation started and ended around the same time may suggest that two processes in the prefrontal and motor regions, respectively, are tightly coupled across time. Future studies should directly address how the two signals interact for optimal motor control.

The neural oscillations results have also identified an interesting post-response decrease in high alpha/low beta power in unexpected compared to expected cue-target alignment and large compared to small target stimulus. These results are in line with previous studies showing alpha and beta band activity related with reward valence and magnitude (Cohen et al., 2008; HajiHosseini and Holroyd, 2015; Mas-Herrero et al., 2015) as well as reward prediction error (Ergo et al., 2019; HajiHosseini et al., 2012). Furthermore, these findings are also in line with the larger literature suggesting increase in cortical processing or neural arousal and increased cortical activity with desynchronization of alpha and beta oscillations (Neuper et al., 2006; Scheeringa et al., 2012). Increased mediofrontal activations, as indicated by the desynchronization in alpha and beta frequency, may imply additional prefrontal processing with regards to the recently completed action. For example, these oscillations could reflect adaptive prefrontal control mechanism that solidifies and promotes the recent course of action that led to larger rewards and an accurate response despite facing an unexpected target.

In summary, our study shows modulation of brain’s motor system by rewards under a real-life context involving dynamic changes where different magnitude of rewards are presented either expectedly or unexpectedly. In particular, by teasing apart the neural processes of movement preparation from execution in the widely investigated LRP signal, we determined the effect of reward magnitude and expectancy across the processing stream of motor hierarchy. In general, our results showed a greater motor preparation when large rewards were more likely and greater motor effort to execute a response when large rewards were presented unexpectedly. These motor activities appeared and ended around the same time as the deployment of prefrontal cognitive control, suggesting a close communication between the two systems to facilitate behavioral performance. Together these results demonstrate an optimized motor control to maximize rewards under the dynamic changes of real-life environment.

